# Effects of Kappa Opioid Receptor Agonists on Fentanyl vs. Food Choice in Male and Female Rats: Contingent vs. Non-Contingent Administration

**DOI:** 10.1101/2020.07.28.225060

**Authors:** E. Andrew Townsend

## Abstract

**Rationale:** Strategies are needed to decrease the abuse liability of mu opioid receptor (MOR) agonists. One strategy under consideration is to combine MOR agonists with kappa opioid receptor (KOR) agonists.

**Objectives:** The effects of KOR-agonists (U50488, nalfurafine) on fentanyl-versus-food choice were compared under conditions where the KOR agonists were added to the self-administered fentanyl (contingent delivery) or administered as pretreatments (non-contingent delivery) in male and female rats. The effects of increasing and decreasing the magnitude of the alternative food reinforcer were also determined.

**Methods:** Rats were trained to respond under a concurrent schedule of fentanyl (0, 0.32-10 μg/kg/infusion) and food reinforcement. In Experiment 1, U50488 and nalfurafine were co-administered with fentanyl as fixed-proportion mixtures (contingent administration). In Experiment 2, U50488 (1-10 mg/kg) and nalfurafine (3.2-32 μg/kg) were administered as acute pretreatments (non-contingent administration). nor-BNI (32 mg/kg) was administered prior to contingent and non-contingent KOR-agonist treatment in Experiment 3. Experiment 4 evaluated the effects of increasing and decreasing the magnitude of the non-drug reinforcer.

**Results:** Both U50488 and nalfurafine decreased fentanyl choice when administered contingently, demonstrating that KOR agonists punish opioid choice. Non-contingent U50488 and nalfurafine administration decreased rates of fentanyl and food self-administration without altering fentanyl choice. Both contingent and non-contingent U50488 and nalfurafine effects on fentanyl choice were attenuated by nor-BNI. Fentanyl choice was sensitive to increases and decreases in the magnitude of the non-drug reinforcer.

**Conclusions:** These results demonstrate that the effects of KOR agonists on fentanyl reinforcement are dependent upon the contingencies under which they are administered.

## 1. Introduction

In 2018, the number of mu opioid receptor (MOR) agonist-related overdose deaths decreased in the United States for the first time in decades, attributable primarily to a decline in fatalities involving prescription opioids over the prior year (Wilson et al. 2020). This welcome reduction in MOR agonist-related overdoses was preceded by a 19% reduction in rates of opioid prescriptions in the United States from 2006 to 2017 (CDC and Prevention 2019), suggesting that decreasing the supply of prescription opioids was an effective approach for reducing the frequency of overdose. However, concerns have been raised that this reduction in prescription opioid availability has led to the inadequate treatment of pain (Pergolizzi et al. 2019). One strategy for addressing the interdependent clinical issues of opioid abuse and pain management includes decreasing the abuse liability of prescription MOR agonists (CDER 2015), with the goal of minimizing the likelihood of misuse while maintaining appropriate access to opioid analgesics to those in pain.

Previous work suggests that kappa opioid receptor (KOR) agonists may decrease the abuse liability of drugs of abuse, including MOR agonists. Stimulation of KORs promotes inhibitory signal transduction through G_i/o_ processes, locally decreasing neuronal excitability and neurotransmitter release (Di Chiara and Imperato 1988; Walker et al. 1987). KORs are highly enriched throughout mesocorticolimbic structures (ventral tegmental area, nucleus accumbens, prefrontal cortex), and KOR agonists inhibit dopamine accumulation within this pathway (Devine et al. 1993; Donzanti et al. 1992; Spanagel et al. 1990). Administration of KOR agonists can also diminish the ability of drugs of abuse to promote mesocorticolimbic dopamine accumulation (e.g., (Margolis et al. 2003; Thompson et al. 2000)), illustrating an opposing interaction between KOR and dopaminergic neurotransmitter systems (for review, see (Escobar et al. 2020; Tejeda and Bonci 2019)).

An implication of this KOR/dopamine interaction is that KOR agonists would be expected to counteract the dopaminergic reinforcing effects of drugs of abuse. Consistent with this hypothesis, KOR agonists decrease rates of cocaine and MOR agonist self-administration under a variety of circumstances (Bowen et al. 2003; Cosgrove and Carroll 2002; Glick et al. 1995; Glick et al. 1998; Mello and Negus 1998; Negus et al. 1997; Negus et al. 2008; Schenk et al. 1999; Schenk et al. 2001; Townsend et al. 2017; Zamarripa et al. 2020) but see (Kuzmin et al. 1997). However, one interpretive complication of this literature is that reinforcement was determined using rate-based schedules of reinforcement. This is an important consideration when evaluating the effects of KOR-agonist effects, because previous works have shown KOR agonists to similarly decrease rates of drug- and food-maintained responding under rate-based schedules of reinforcement (Cosgrove and Carroll 2002; Mello and Negus 1998; Negus et al. 1997; Negus et al. 2008). Thus, the aforementioned studies do not exclude the possibility that KOR agonists decrease rates of drug self-administration through non-selective effects on operant responding.

One method to minimize the influence of non-selective effects on operant responding includes the use of concurrent “choice” schedules of reinforcement. The primary dependent measure under concurrent schedules is behavioral allocation, which has been shown to be insensitive to alterations in response rate (for review, see (Banks and Negus 2012)). Three studies have utilized choice procedures to evaluate KOR-agonist effects on drug reinforcement. In the earliest study, acute pretreatment with the KOR agonist enadoline failed to attenuate cocaine-versus-money choice in humans (Walsh et al. 2001a). A later study using rhesus monkeys as subjects found continuous 3-day treatment with the KOR agonist U50488 to increase the choice of cocaine over a food alternative (Negus 2004). These findings do not support the hypothesis that KOR agonists selectively decrease the reinforcing effects of drugs of abuse, with the latter results suggesting that KOR-agonist maintenance may actually enhance the reinforcing effects of cocaine. However, a more recent study suggests the experimental conditions under which a KOR agonist is administered is an important determinant of its effects on drug reinforcement. Using a drug-versus-drug choice procedure in rhesus monkeys, the KOR agonist salvinorin A decreased drug + salvinorin A choice, irrespective of whether the reinforcer was remifentanil or cocaine (Freeman et al. 2014). This finding suggests that *contingent* KOR-agonist administration (i.e., operant responding results in KOR agonist delivery) punishes drug self-administration, as the introduction of the KOR agonist as a consequent stimulus decreased the probability that drug self-administration would occur (definition of punishment: (Azrin 1966)).

A translational implication of the aforementioned results is that a KOR agonist would only be expected to selectively decrease the reinforcing effects of a MOR agonist if it were administered contingently, which could be accomplished by combining a KOR agonist and a MOR agonist into a single medication. However, previous reports of poor tolerability of KOR agonists introduces a potential obstacle to the clinical utility of this approach. Namely, dysphoric, psychotomimetic, and sedative effects have been reported following acute KOR agonist administration in humans (Kumor et al. 1986; Pfeiffer et al. 1986; Walsh et al. 2001b). These unpleasant KOR-agonist effects could contribute to medication non-compliance, resulting in inadequate pain management.

In recent years, a new class of *G-protein biased* KOR agonists have emerged, and evidence suggests that these compounds may produce fewer adverse effects than traditional KOR agonists (Mores et al. 2019). Relative to traditional KOR agonists that activate intracellular G-protein and beta-arrestin pathways with similar potency (e.g., U50488), G-protein biased KOR agonists exhibit greater potency to activate the G-protein pathway. In light of previous work that supports a role of beta-arrestin activation in the aversive effects of KOR agonists (Bruchas and Chavkin 2010; Bruchas et al. 2007), G-protein biased KOR agonists may produce fewer untoward effects and have a greater likelihood of tolerability. One of these drugs is nalfurafine, which functions as a G-protein-biased agonist at both human and rat KOR (Kaski et al. 2019; Liu et al. 2019; Schattauer et al. 2017). Nalfurafine is the only selective KOR agonist approved for clinical usage, and it has been available for the treatment of uremic pruritus in Japan since 2009. No clinical reports of psychiatric side effects have been reported (Kozono et al. 2018), providing evidence that nalfurafine is well tolerated. Recent preclinical studies have reported that contingently administered nalfurafine can punish oxycodone self-administration in rats and rhesus monkeys when the two drugs are self-administered as a mixture (Townsend et al. 2017; Zamarripa et al. 2020). While encouraging, these studies used rate-based schedules to evaluate oxycodone reinforcement. Therefore, it remains unclear whether nalfurafine decreased rates of MOR agonist self-administration through a non-selective effect other than punishment.

The current study compared the effects of contingent and non-contingent KOR agonist administration on fentanyl-versus-food choice in male and female rats. We hypothesized that contingent KOR-agonist administration would punish opioid choice, and non-contingent KOR-agonist pretreatment would fail to selectively affect opioid self-administration. The current study also sought to evaluate whether a G-protein biased KOR agonist (nalfurafine) produced similar effects on fentanyl-versus-food choice relative to an unbiased compound (U50488). The effects of contingent and non-contingent KOR agonist administration were compared to those of the non-pharmacological intervention of increasing and decreasing the magnitude of the alternative food reinforcer.

## 2. Methods

### 2.1. Subjects

Twenty-three Sprague-Dawley rats (11 male, 12 female) were acquired at 10 weeks of age (Envigo Laboratories, Frederick, MD, USA) and surgically implanted with vascular access ports (Instech, Plymouth Meeting, PA) with custom-made jugular catheters as described previously (Huskinson et al. 2017). Rats were divided into two cohorts. Cohort 1 (5 male, 6 female) was used in Experiments 1, 2, and 3. Cohort 2 (6 male, 6 female) was used in Experiment 4. Although all twelve rats of Cohort 2 completed Experiment 4, only eight of the eleven rats of Cohort 1 completed Experiments 2 and 3 (3 male, 5 female). Rats were singly housed in a temperature and humidity-controlled vivarium, maintained on a 12-h light/dark cycle (lights off at 6:00 PM). Water and food (Teklad Rat Diet, Envigo) were provided ad-libitum in the home cage. Behavioral testing was conducted five days per week from approximately 2:00 PM - 4:00 PM. Catheter patency was verified at the conclusion of each Experiment by instantaneous muscle tone loss following intravenous (IV) methohexital (0.5 mg) administration. Animal maintenance and research were conducted in accordance with the 2011 guidelines for the care and use of laboratory animals and protocols were approved by the Virginia Commonwealth University Institutional Animal Care and Use Committee.

### 2.2. Apparatus and Catheter Maintenance

Twelve modular operant chambers located in sound-attenuating cubicles (Med Associates, St. Albans, VT, USA) were equipped with two retractable levers, a set of three LED lights (red, yellow, green) mounted above each lever, and a retractable “dipper” cup (0.1 ml) located between the levers for presenting diluted Ensure® (0, 1.8, 18 or 100% v/v vanilla flavor Ensure® in tap water; Abbott Laboratories, Chicago, IL, USA). Intravenous drug solutions were delivered by activation of a syringe pump (PHM-100, Med Associates) located inside the sound-attenuating cubicle as described previously (Townsend et al. 2017). After each behavioral session, catheters were flushed with gentamicin (0.4 mg), followed by 0.1 ml of heparinized saline (10 U/ml).

### 2.3. Drugs

Fentanyl HCl and nalfurafine HCl were provided by the National Institute on Drug Abuse (NIDA) Drug Supply Program (Bethesda, MD) and dissolved in sterile saline. (±) U50488 HCl was purchased commercially (Tocris, Pittsburgh, PA) and dissolved in sterile saline. Nor-BNI di-HCl (synthesized by K Cheng and K Rice, National Institutes of Health, Bethesda, MD) was dissolved in 1 percent lactic acid : sterile water at a concentration of 22 mg/ml. Methohexital was purchased from the Virginia Commonwealth University pharmacy (Richmond, VA). All solutions were passed through a 0.22-micron sterile filter (Millex GV, Millipore Sigma, Burlington, MA) before administration. All drug doses were expressed as the salt forms listed above and delivered based on weights collected weekly.

### 2.4. Procedure

#### 2.4.1. Self-administration Training

Rats were trained to respond in a fentanyl-versus-food choice procedure as described previously (Townsend et al. 2019a; Townsend et al. 2019b). Briefly, rats were first trained to respond on the right lever for IV fentanyl (3.2 μg/kg/infusion) under a fixed-ratio 5, 20s timeout (FR5, TO20) schedule of reinforcement, signaled by the illumination of a green stimulus light. Next, rats were trained to respond on the left lever for a 5s presentation of 18% Ensure® under a FR5, TO20 schedule of reinforcement, signaled by the illumination of a red stimulus light. Once rats were trained to respond for fentanyl and 18% Ensure® in isolation, both reinforcers were made available under a concurrent FR5, TO20: FR5, TO20 schedule of reinforcement. Here, the behavioral session consisted of five 20-min response components each preceded by a 4-min “sample” component. Each sample component started with a non-contingent infusion of the unit fentanyl dose available during the upcoming response component followed by a 2-min time out. Next, a 5-s presentation of liquid food was programmed followed by a 2-min time out. Following this second time out, the response component began. During each response component, both levers were extended, a red stimulus light above the left lever was illuminated to signal liquid food availability and a green stimulus light above the right lever was illuminated to signal IV fentanyl availability. Response requirement (FR5) completion on the left lever resulted in a 5-s presentation of liquid food whereas response requirement (FR5) completion on the right lever resulted in the delivery of the IV fentanyl dose available for that component. Responding on one lever reset the ratio requirement for the other lever. The Ensure® concentration was held constant throughout the session. A different fentanyl dose was available during each of the five successive response components (0, 0.32, 1.0, 3.2, and 10 μg/kg/inf during components 1–5, respectively). Fentanyl dose was varied by changing the infusion duration (e.g., 315 g rat: 0, 0.5, 1.56, 5, and 15.6 s during components 1–5, respectively) and the green light above the fentanyl-lever flashed on and off in 3-s cycles (i.e., longer flashes corresponded with larger fentanyl doses).

During each response component, rats could complete up to 10 total ratio requirements between the food- and fentanyl-associated levers. Each ratio requirement completion initiated a 20 s time out, the retraction of both levers, and extinction of the red and green stimulus lights. If all 10 ratio requirements were completed before 20 min had elapsed, then both levers retracted, and stimulus lights were extinguished for the remainder of that component. Choice was considered stable when the smallest fentanyl dose that maintained at least 80% of completed ratio requirements on the fentanyl-associated lever was within a 0.5 log unit of the running mean for three consecutive days with no trends (i.e., stability criteria).

#### 2.4.2. Experiment 1: Effects of Contingent U50488 and Nalfurafine Administration on Fentanyl vs Food Choice

Once the fentanyl-vs.-food-training criteria were met, the punishing effects of contingently administered U50488 and nalfurafine were evaluated in Cohort 1 rats. Each fentanyl/KOR agonist mixture was tested for at least 5 sessions and until stability criteria were met. All fentanyl:nalfurafine mixtures were tested in a counterbalanced dosing order (fentanyl:nalfurafine: 1:0 (fentanyl alone), 1:0.1, 1:0.32, 1:1). Fentanyl:U50488 mixtures were similarly tested in a counterbalanced dosing order (fentanyl:U50488: 1:0 (fentanyl alone), 1:10, 1:32, 1:100).

#### 2.4.3. Experiment 2: Effects of Non-contingent U50488 and Nalfurafine Administration on Fentanyl vs Food Choice

The effects of non-contingent KOR agonist administration on fentanyl-vs.-food choice were also evaluated in Cohort 1 rats. Rats received a subcutaneous injection of KOR agonist or vehicle (saline) 10 min prior to every other choice session, with a fentanyl-vs.-food choice session in the absence of KOR-agonist pretreatment separating each test condition. Nalfurafine (saline, 3.2, 10, and 32 μg/kg) and U504488 (saline, 1, 3.2, 10 mg/kg) pretreatments were tested in a counterbalanced order. Nalfurafine was tested before U50488. Pretreatment times were based on previous U50488 and nalfurafine intracranial self-stimulation studies (Faunce and Banks 2020; Lazenka et al. 2018).

#### 2.4.4. Experiment 3: Effects of nor-BNI on U50488 and Nalfurafine Administration on Fentanyl vs Food Choice

The effects of nor-BNI (selective KOR antagonist) treatment on both contingent and non-contingent U50488 and nalfurafine administration were determined as a final experiment in Cohort 1. A single intraperitoneal injection of nor-BNI (32 mg/kg, i.p.) occurred 24 h prior to behavioral studies, accounting for the slow onset and long duration of action of the compound (Endoh et al. 1992). nor-BNI effects were evaluated across 5 daily testing sessions, wherein 1) KOR-agonist vehicle (saline), 2) 1:100 (fentanyl:U50488), 3) 1:1 (fentanyl:nalfurafine), 4) 10 mg/kg U50488, and 5) 32 μg/kg nalfurafine were tested in a counterbalanced order. Pretreatment times for non-contingent KOR-agonist injections were 10 min.

#### 2.4.5. Experiment 4: Effects of Food Reinforcer Manipulation on Fentanyl vs Food Choice

Once the aforementioned fentanyl-vs.-food-training criteria were met using 18% diluted Ensure® as the food reinforcer, the effects of increasing and decreasing the Ensure® concentration were evaluated in Cohort 2 rats. Ensure® concentrations (1.8, 18, 100%) and water (0%) were tested for two consecutive sessions in a counterbalanced order.

### 2.5. Data Analysis

The two primary dependent measures were 1) percent drug choice, defined as (number of ratio requirements, or ‘choices’, completed on the fentanyl-associated lever/total number of choices completed on both levers) × 100, and 2) number of choices per component. For Experiment 1, data from the last three stable testing sessions were averaged for analysis. For Experiments 2 and 3, analysis was performed from the single test session of each condition. For Experiment 4, each condition was in effect for two days, with the data from the second day used for data analysis. Results were plotted as a function of unit fentanyl dose and analyzed using a two-way ANOVA or a mixed-model analysis in instances of missing data (noted in figures) (GraphPad Prism 8, La Jolla, CA). Baseline choice functions were re-determined for each Experiment and KOR agonist. These five baseline functions were compared within Cohort 1 using a mixed-model analysis. For each Experiment, data were separated by sex and analyzed using a distinct two-way ANOVA or mixed-model analysis for each experimental condition. For these sex comparisons, the *p* values of related tests were corrected for Type I error using the Benjamini and Hochberg procedure (Keselman et al. 2002) within each Experiment. In the presence of a significant main effect or interaction, a Dunnett post-hoc test was conducted as appropriate. Statistical significance was set a priori at the 95% confidence level (*p*<0.05).

## 3. Results

### 3.1. Baseline Fentanyl vs food choice

Under baseline conditions, liquid food was almost exclusively chosen when no fentanyl or the smallest unit dose of fentanyl (0.32 μg/kg/infusion) was available (dashed lines; **Figure 1A, 1B**, **2A, 2B**, **3A**, **4A**). As the fentanyl dose increased, behavior was reallocated to the fentanyl lever and the largest fentanyl dose (10 μg/kg/infusion) maintained near exclusive fentanyl choice. Additionally, choices per component decreased as a function of increasing fentanyl doses (dashed lines; **Figure 1C, 1D**, **2C, 2D**, **3B**, **4B**). Baseline choice behavior remained stable across Experiments 1, 2, and 3 in Cohort 1 (*percent fentanyl choice*: fentanyl dose: F_1.2, 10.9_ = 103.5, *p*<0.0001; Experiment: F_1.6, 14.4_ = 1, *p*=0.36; interaction: F_3.4, 23.1_ = 1.2, *p*=0.32; *choices per component*: fentanyl dose: F_2.7, 24.3_ = 230.4, *p*<0.0001; Experiment: F_2.5, 22.5_ = 1.2, *p*=0.31; interaction: F_3.3, 23.3_ = 0.75, *p*=0.54). No effect of sex was detected on either dependent measure under baseline or vehicle-treatment conditions.

**Fig 1.**
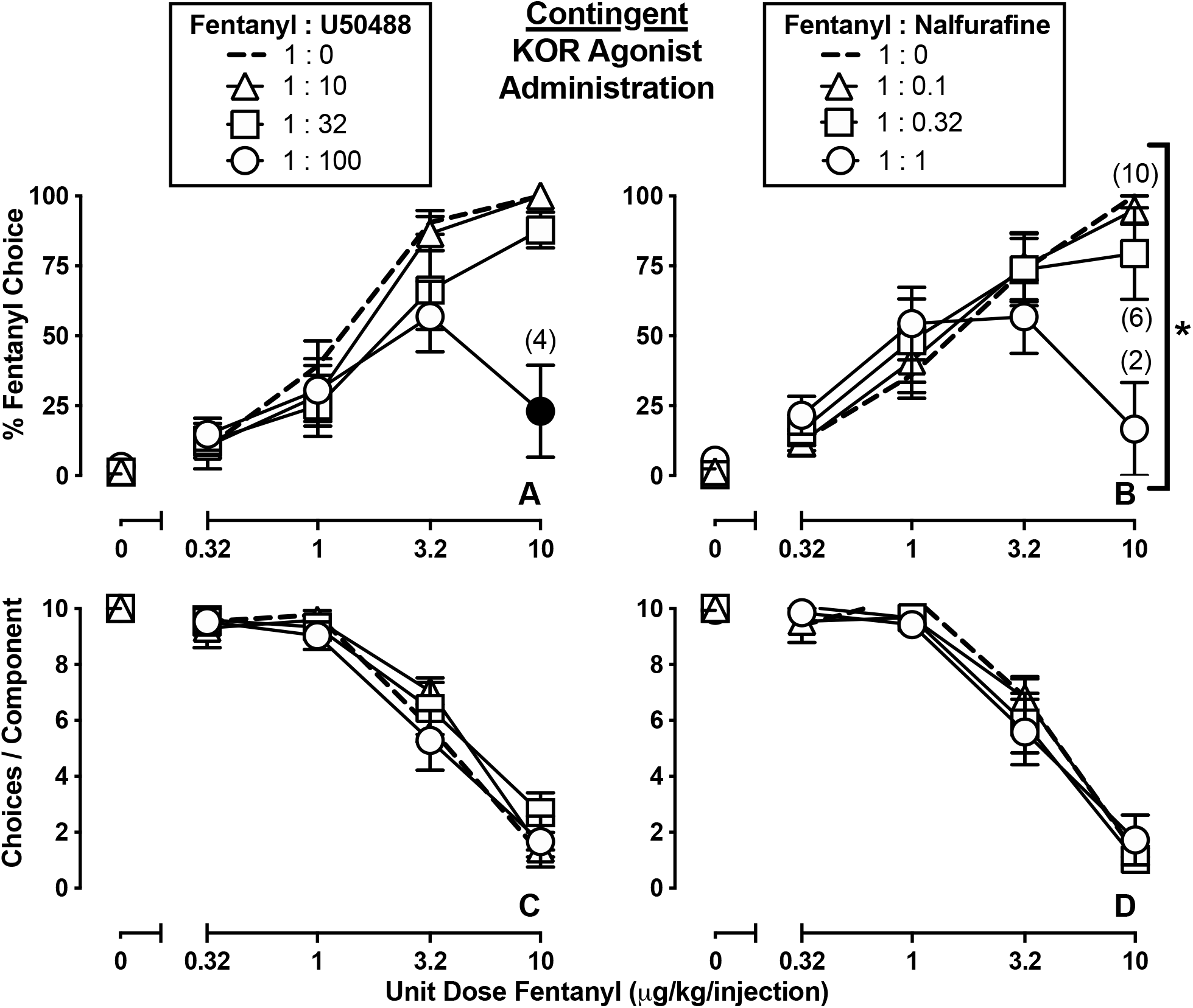
Effectiveness of KOR agonists to punish choice of fentanyl over food in male and female rats. Abscissa: IV fentanyl unit dose in μg/kg. Top row ordinates: Percentage of completed ratio requirements on the fentanyl-associated lever. Bottom row ordinates: Number of choices completed per component. Left Panels: Effects of contingently administered U50488 on percent fentanyl choice (**A**) and the number of choices completed per component (**C**) (n=6 female, 4 male). Right Panels: Effects of contingently administered nalfurafine on percent fentanyl choice (**B**) and the number of choices completed per component (**D**) (n=6 female, 5 male). Points represent mean ± SEM, numbers in parenthesis denote the number of rats that completed at least one ratio requirement at a given data point in instances wherein a subset of rats did not respond, and filled symbols denote significant difference relative to baseline. * denotes a significant fentanyl unit dose × fentanyl:nalfurafine mixture proportion interaction. Significance defined as *p* < 0.05.

### 3.2. Experiment 1: Effects of Contingent U50488 and Nalfurafine Administration on Fentanyl vs Food Choice

The unbiased KOR agonist, U50488, functioned as a punisher of fentanyl choice (**Figure 1A**: fentanyl dose: F_2, 17.9_ = 0.5, *p*<0.0001; fentanyl:U50488 mixture proportion: F_2.3, 20.7_ = 0.76, *p*=0.003; interaction: F_5, 40_ = 0.41, *p*=0.0007), with the 1:100 (fentanyl:U50488) proportion decreasing choice of the largest fentanyl unit dose. Although a unit fentanyl dose and fentanyl/nalfurafine mixture interaction was detected (**Figure 1B**: fentanyl dose: F_1.7, 17.2_ = 0.43, *p*<0.0001; fentanyl:nalfurafine mixture proportion: F_1.8, 18_ = 2.5, *p*=0.12; interaction: F_2.4, 20_ = 4.8, *p*=0.016), post-hoc analysis did not detect significant changes in fentanyl choice relative to fentanyl alone. Although some rats did not respond when the largest unit dose of fentanyl was combined with U50488 or nalfurafine, the average number of choices completed per component was not significantly affected by contingent KOR agonist administration (**Figure 1C** and **1D**, respectively). The number of choices completed per component was lower in male rats relative to female rats when the 1:100 (fentanyl:U50488) proportion was available (adjusted *p* = 0.04). No other effect of sex was detected on either dependent measure following contingent administration of U50488 or nalfurafine.

### 3.3. Experiment 2: Effects of Non-contingent U50488 and Nalfurafine Administration on Fentanyl vs Food Choice

Pretreatment with U50488 or nalfurafine did not significantly alter behavioral allocation between fentanyl and food (U50488: **Figure 2A**; nalfurafine: **Figure 2B**). However, both KOR agonists decreased the number of choices completed per component (*U50488*: **Figure 2C**: fentanyl dose: F_2.8, 19.6_ = 39.9, *p*<0.0001; fentanyl:U50488 mixture proportion: F_1.8, 12.6_ = 31, *p*<0.0001; interaction: F_2.9, 20.4_ = 6.6, *p*=0.003; *nalfurafine*: **Figure 2D**: fentanyl dose: F_1.7, 11.6_ = 41.9, *p*<0.0001; fentanyl:nalfurafine mixture proportion: F_2.2, 15.2_ = 10.9, *p*=0.001; interaction: F_3.8, 26.5_= 4, *p*=0.013). Fentanyl choice was greater in male rats relative to female rats following pretreatment with 3.2 μg/kg nalfurafine (adjusted *p* = 0.03). No other effect of sex was detected on either dependent measure following non-contingent U50488 or nalfurafine administration.

**Fig 2.**
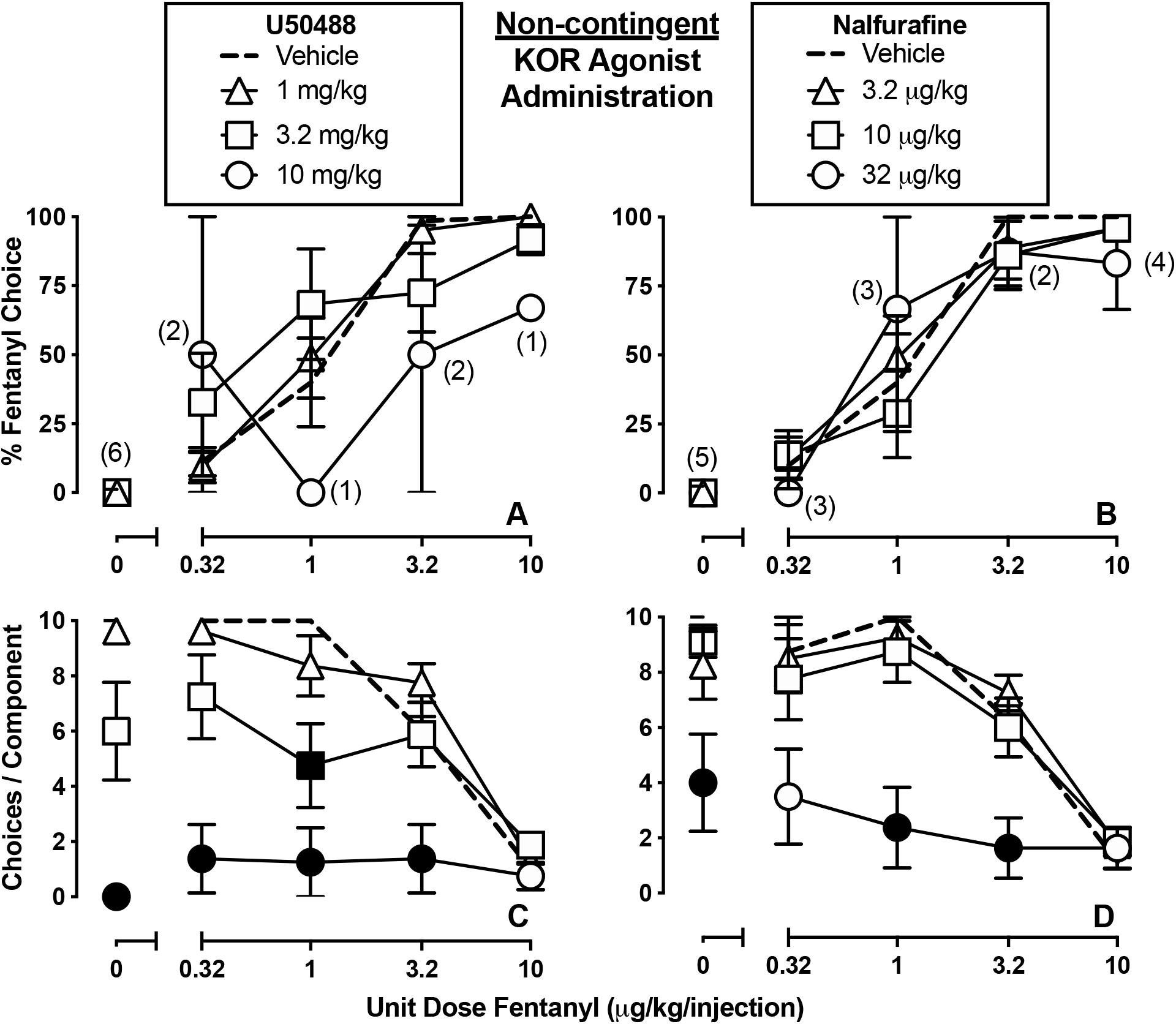
Effectiveness of non-contingent administration of KOR agonists on fentanyl-vs.-food choice in male and female rats. Abscissa: IV fentanyl unit dose in μg/kg. Top row ordinates: Percentage of completed ratio requirements on the fentanyl-associated lever. Bottom row ordinates: Number of choices completed per component. Left Panels: Effects of non-contingent U50488 pretreatment on percent fentanyl choice (**A**) and the number of choices completed per component (**C**) (n=5 female, 3 male). Right Panels: Effects of non-contingent nalfurafine pretreatment on percent fentanyl choice (**B**) and the number of choices completed per component (**D**) (n=5 female, 3 male). Points represent mean ± SEM, numbers in parenthesis denote the number of rats that completed at least one ratio requirement at a given data point in instances wherein a subset of rats did not respond, and filled symbols denote significant difference relative to baseline. Of note, none of the rats responded during the 0 μg/kg/injection component following injection of 10 mg/kg U50488. Significance defined as *p* < 0.05.

### 3.4. Experiment 3: Effects of nor-BNI on U50488 and Nalfurafine Administration on Fentanyl vs Food Choice

Administration of the KOR antagonist nor-BNI blocked the effects of contingent and non-contingent KOR agonist administration on fentanyl choice and the number of choices completed per component (**Figure 3**). An effect of sex was detected for fentanyl choice of the 1:1 (fentanyl:nalfurafine) mixture following nor-BNI treatment (adjusted *p* = 0.011), with increased fentanyl choice in male rats. No other effect of sex was detected on either dependent measure in Experiment 3.

**Fig 3.**
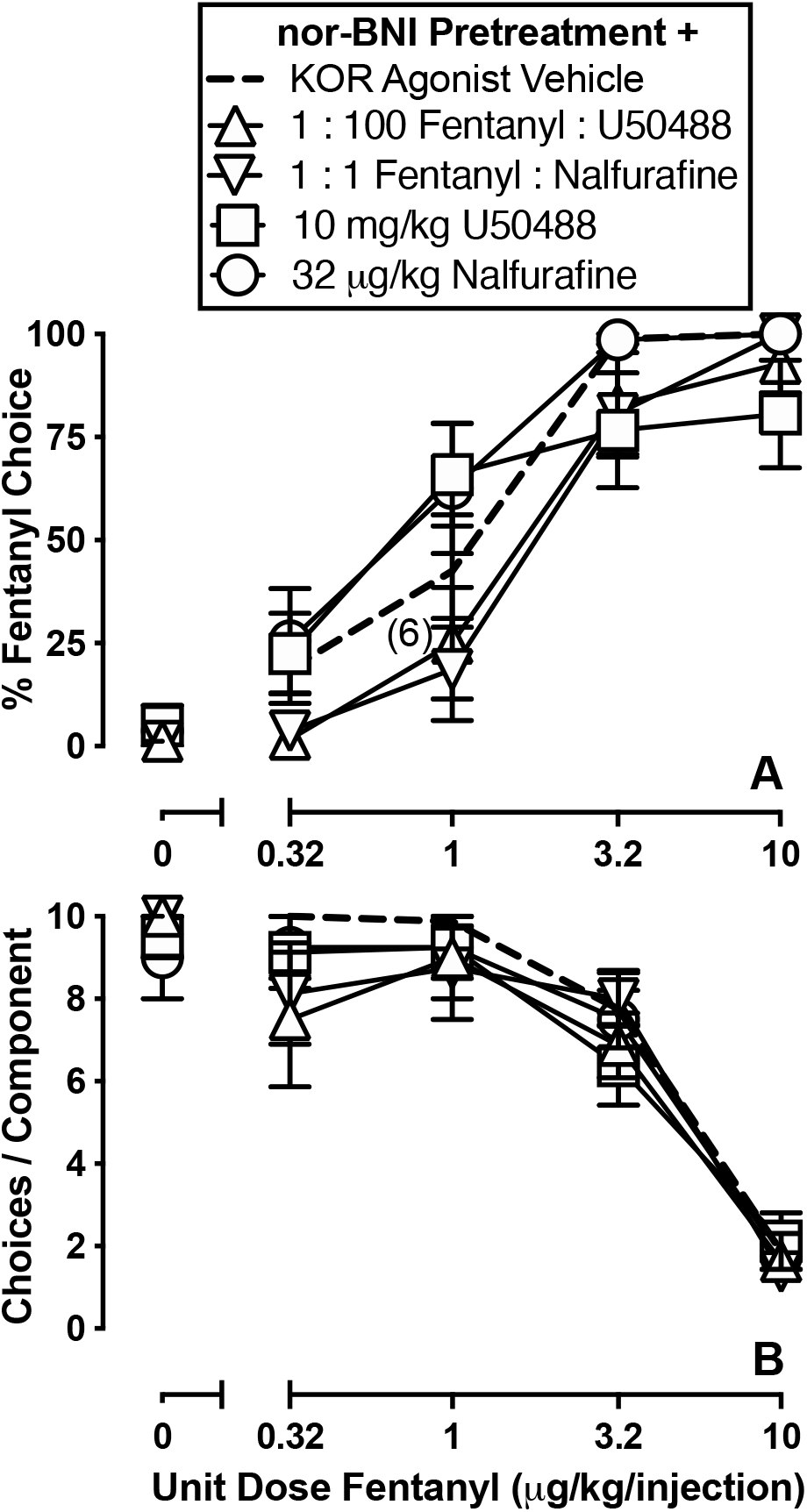
Effectiveness of nor-BNI to block the effects of contingent and non-contingent KOR-agonist administration on fentanyl-vs.-food choice in male and female rats (n=5 female, 3 male). Abscissa: IV fentanyl unit dose in μg/kg. Top row ordinate (**A**): Percentage of completed ratio requirements on the fentanyl-associated lever. Bottom row ordinate (**B**): Number of choices completed per component. Points represent mean ± SEM, numbers in parenthesis denote the number of rats that completed at least one ratio requirement at a given data point in instances wherein a subset of rats did not respond (i.e., 1:100 fentanyl:U50488), and filled symbols denote significant difference relative to baseline. Significance defined as *p* < 0.05.

### 3.5. Experiment 4: Effects of Food Reinforcer Manipulation on Fentanyl vs Food Choice

Fentanyl choice was sensitive to both increases and decreases in the magnitude of the alternative reinforcer (**Figure 4A**: fentanyl dose: F_2.2, 23.7_ = 166, *p*<0.0001; Ensure® concentration: F_2.6, 28.2_ = 49.2, *p*<0.0001; interaction: F_4.2, 45.8_ = 6.5, *p*=0.0003). Relative to 18% Ensure® (i.e., concentration used in Experiments 1, 2, and 3), percent choice of 0, 0.32, and 1 μg/kg/infusion fentanyl decreased when the alternative food reinforcer was 100% (undiluted) Ensure. In contrast, percent choice of 0.32, and 1 μg/kg/infusion fentanyl was increased when the alternative reinforcer was water (0%) or highly diluted Ensure® (1.8%). Manipulation of the alternative reinforcer also affected the number of choices completed per component (**Figure 4B**: fentanyl dose: F_2, 22.4_ = 233.4, *p*<0.0001; Ensure® concentration: F_1.7, 19.2_ = 11.1, *p*=0.0009; interaction: F_3.8, 41.8_ = 2.5, *p*=0.06). No effect of sex was detected on either dependent measure in Experiment 4.

**Fig 4.**
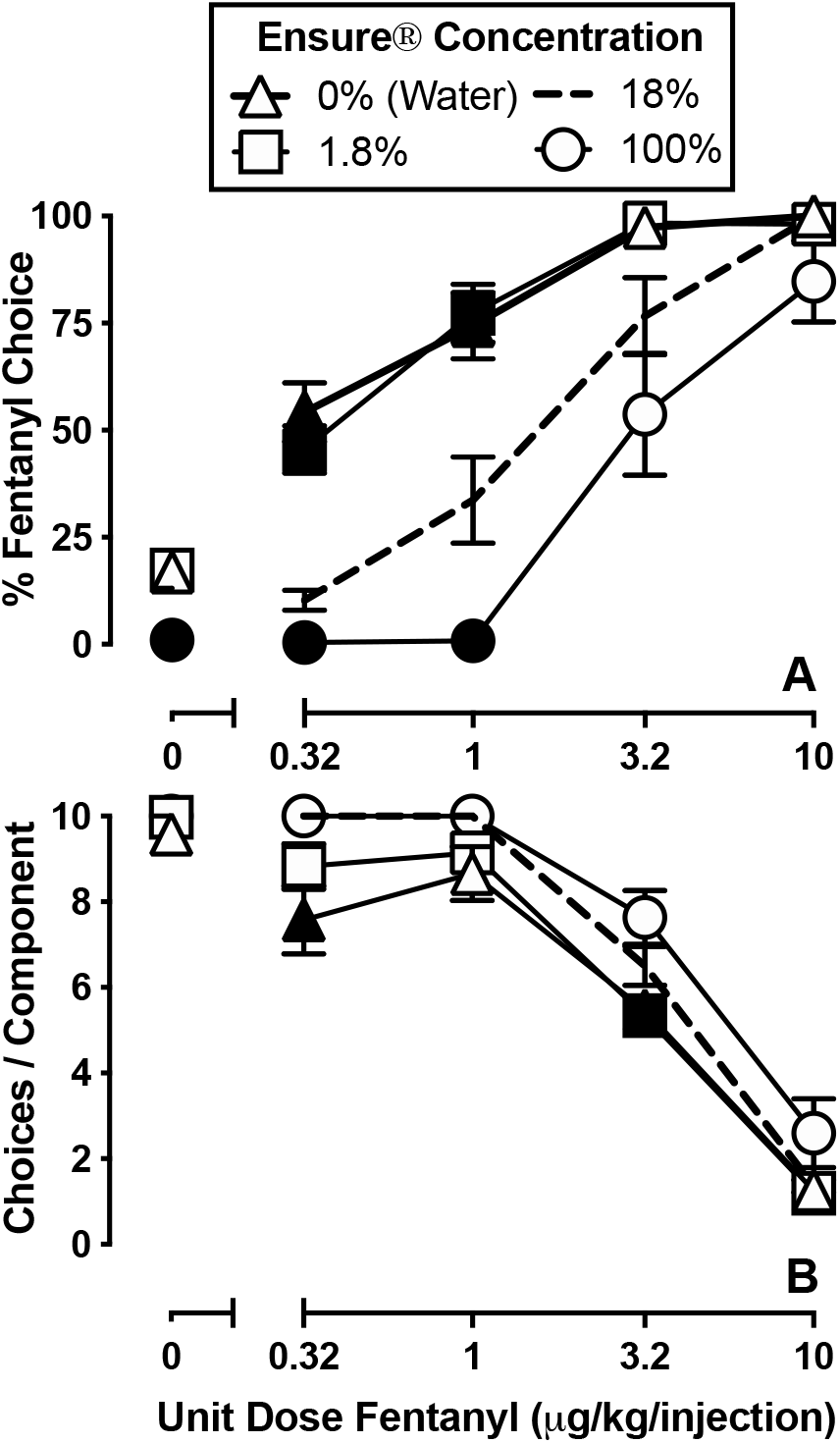
Effects of food reinforcer magnitude on fentanyl-vs.-food choice in male and female rats (n=6 female, 6 male). Abscissa: IV fentanyl unit dose in μg/kg. Top row ordinate (**A**): Percentage of completed ratio requirements on the fentanyl-associated lever. Bottom row ordinate (**B**): Number of choices completed per component. Points represent mean ± SEM and filled symbols denote significant difference relative to baseline. Significance defined as *p* < 0.05.

## 4. Discussion

The current study demonstrated that KOR-agonists punished fentanyl self-administration under a choice procedure when administered contingently (i.e. fixed-proportion fentanyl/KOR agonist). In addition, KOR agonists did not significantly attenuate fentanyl self-administration when administered non-contingently up to KOR agonist doses that decreased rates of operant responding. This demonstration of KOR-mediated punishment under an opioid-vs-food choice procedure is in agreement with previous results of salvinorin A punishing cocaine and remifentanil choice (Freeman et al. 2014), and extends the generality of this finding to include the KOR agonists U50488 and nalfurafine. When considered alongside previous reports of nalfurafine’s tolerability in clinical populations (Kozono et al. 2018), nalfurafine-mediated punishment of fentanyl self-administration may be encouraging, as nalfurafine is clinically available and could be repurposed to decrease the abuse liability of opioid analgesics. However, nalfurafine proportions that punished fentanyl choice did so at doses that also produced rate-decreasing effects. The latter results suggest a narrow therapeutic window for KOR-induced punishment vs. undesirable effects.

The putative G-protein biased KOR agonist nalfurafine and the unbiased KOR agonist U50488 similarly punished fentanyl self-administration. Although weaker evidence of punishment was observed with nalfurafine based on statistical analysis, the overall findings were consistent with a recent report of contingently administered nalfurafine and salvinorin A decreasing rates of oxycodone self-administration under a progressive-ratio schedule of reinforcement in rhesus monkeys (Zamarripa et al. 2020). Considered alongside reports that nalfurafine and other G-protein biased KOR agonists attenuate the conditioned rewarding effects of MOR agonists (Kaski et al. 2019; Tsuji et al. 2001), these results suggest that the abuse-limiting effects of KOR agonists may be G-protein mediated. Furthermore, in light of evidence that the aversive effects of KOR agonists are beta-arrestin mediated (Bruchas and Chavkin 2010; Bruchas et al. 2007), these results also suggest that aversion may not be a necessary component of KOR-mediated punishment. However, given that nalfurafine is only moderately G-protein biased, particularly at rat KOR (Schattauer et al. 2017), higher doses of nalfurafine may have recruited beta-arrestin-mediated processes. Therefore, the evaluation of additional KOR agonists with greater intracellular bias for the G-protein pathway (e.g., (Mores et al. 2019)) as well as beta-arrestin biased KOR agonists (e.g., (Crowley et al. 2020)) could clarify the role of G-protein and beta-arrestin signaling in KOR-mediated punishment.

Acute, non-contingent administration of U50488 or nalfurafine failed to selectively decrease fentanyl choice and decreased overall rates of operant behavior. These findings suggest KOR agonists would not be effective as standalone treatments for substance use disorders (see (Banks 2020)) and are in agreement with a report of acute enadoline treatment failing to selectively affect cocaine-versus-money choice in human volunteers (Walsh et al. 2001a). Previous works have highlighted the importance of evaluating behavioral selectivity of treatment effects of candidate substance use disorder treatments (Czoty et al. 2016; Haney and Spealman 2008; Mello and Negus 1996), arguing that such measures increase the potential for preclinical-to-clinical translation. Collectively, these results illustrate the utility of concurrent schedules of reinforcement for dissociating the punishing and non-selective rate-decreasing effects of KOR agonists.

The choice of smaller doses of fentanyl was decreased when the magnitude of the alternative food reinforcer (Ensure®) was increased from 18 to 100%, and the opposite was observed when the Ensure® concentration was decreased to 1.8%. Nevertheless, choice of the largest dose of fentanyl was unaffected by Ensure® manipulation, thus monotonic increases in fentanyl choice were maintained despite manipulations of the alternative reinforcer. These results are consistent with previous reports of manipulating the magnitude of the non-drug reinforcer under similar concurrent schedules of reinforcement (e.g., (Negus 2003; Thomsen et al. 2013; Townsend et al. 2019b)). Findings of surmountability have also been reported in evaluations of the effectiveness of electric shock (Johanson 1975; Johanson 1977) and histamine (Negus 2005) to punish cocaine choice, suggesting that the punishing effectiveness of a given stimulus decreases as the dose of the drug reinforcer increases. In contrast, the current report found the punishing effectiveness of U50488 and nalfurafine to be most evident when the largest dose of fentanyl (10 μg/kg/infusion) was available. Differences in the delivery of the punishing stimuli may account for these differences. In the previous studies, the punishing effectiveness of a fixed intensity of electric shock (Johanson 1975; Johanson 1977) or a fixed dose of histamine (Negus 2005) was evaluated across a range of cocaine doses. In the current study, KOR agonists were administered as a fixed mixture proportion with fentanyl, wherein the doses of each drug were increased in tandem. An implication of the current results is that formulating an opioid analgesic with a pharmacological punisher may limit the reinforcing dose range of the opioid analgesic, as consumption of higher doses of the opioid analgesic would be met with higher doses of the pharmacological punisher. However, the proportion of opioid analgesic to punisher would need to be carefully considered, with the amount of punisher being sufficient to discourage overconsumption while being low enough such that appropriate clinical usage is unpunished, else risking medication non-compliance.

Although evidence for punishment was detected with both KOR agonists, it also corresponded with decreases in operant behavior in some rats. This may reflect KOR-mediated sedation, as a recent study found that combining nalfurafine or U50488 with oxycodone enhanced sedation-like effects in rhesus monkeys (Huskinson et al. 2020). However, foundational work evaluating the effectiveness of electric shock to punish food-maintained responding noted that rates of responding decrease following the delivery of electric shock (Azrin 1959; Azrin 1966; Bergman and Johanson 1981; Grove and Schuster 1974). Given that electric shock does not produce sedative effects, these findings suggest rates of behavior decrease following delivery of punishing stimuli and that the rate-decreasing effects observed in the current study may not necessarily reflect sedation. Future studies could increase the session duration or use a discrete trial choice procedure to limit potential carryover effects from the previous self-administered drug or drug mixture. In addition, preclinical “drug plus food” choice procedures could evaluate whether mixtures of MOR and KOR agonists 1) lack reinforcing effects or 2) serve as punishing stimuli themselves (e.g., (Minervini et al. 2019)). Ultimately, human-laboratory studies are needed to evaluate the relationship between the reinforcing, punishing, and subjective effects of opioid analgesic/KOR agonist mixtures.

No sex differences in the punishing effectiveness of U50488 or nalfurafine were detected. However, the rate-decreasing effects of the 1:100 (fentanyl:U5088) mixture were greater in male rats. These data are consistent with previous reports of male rodents being more sensitive to the locomotor-suppressant effects of U50488 (Craft and Bernal 2001; Kavaliers and Innes 1987). Male rats were also found to choose more fentanyl than female subjects following non-contingent administration of the smallest tested dose of nalfurafine (3.2 μg/kg), although these effects were not detected at larger nalfurafine doses or any tested dose of U50488. Male rats also exhibited increased choice of the 1:1 (fentanyl:nalfurafine) mixture following treatment with nor-BNI. The effects of KOR agonist treatment were quantitatively similar between sexes otherwise. Nevertheless, subtle sex differences in KOR agonist effects were detected. In light of previous reports of increased sensitivity to the punishing effects of electric shock in female rodents (Kutlu et al. 2020; Orsini et al. 2016), these results provide rationale for the further evaluation of the interaction between sex as a biological variable and punishment.

The current results illustrate that the effects of a drug on behavior can be more complex than its interactions with receptors. Experimenter-administered delivery of KOR agonists to the subject was found to produce non-specific decreases in rates of both fentanyl and Ensure® reinforcement. However, if delivery of the KOR agonist was the consequence of fentanyl self-administration by the subject, the relative reinforcing effects of fentanyl were decreased. A potential clinical implication of this finding is that formulating a KOR agonist with an opioid analgesic (MOR agonist) may decrease the likelihood of misuse and overconsumption. An additional and intriguing question for future research is whether the contingency-dependent effects of KOR agonists can be dissociated from a neurochemical, circuit, or other quantitative neurobiological measure. Recent efforts have addressed similar questions related to the punishing effects of electric shock (e.g., (Jacobs and Moghaddam 2020; Kutlu et al. 2020; Verharen et al. 2020)). Expanding this area of research to include pharmacological punishers such as KOR agonists could elucidate both similarities and differences between KOR- and shock-mediated punishment, which may aid in the identification of substrates that mediate environmental and contingency-dependent determinants of decision making.

## 5. Acknowledgements

This work was supported by 1) a Research Grant awarded by the Virginia Commonwealth University Postdoctoral Association and 2) the National Institute on Drug Abuse of the National Institutes of Health under Award Numbers F32DA047026 and P30DA033934, and 3) the National Institute on Alcohol Abuse and Alcoholism and the National Institute on Drug Abuse Intramural Research Programs. The manuscript content is solely the responsibility of the author and does not necessarily reflect the official views of the National Institutes of Health. The author would like to thank Dr. Matthew L. Banks, Dr. Kevin B. Freeman, and Mr. C. Austin Zamarripa for comments on an earlier version of the manuscript.

## Notes

No conflicts of interest

Funding and Disclosure: This work was supported by 1) a Research Grant awarded by the Virginia Commonwealth University Postdoctoral Association, 2) the National Institute on Drug Abuse of the National Institutes of Health under Award Numbers F32DA047026 and P30DA033934, and 3) the National Institute on Alcohol Abuse and Alcoholism and the National Institute on Drug Abuse Intramural Research Programs. The manuscript content is solely the responsibility of the author and does not necessarily reflect the official views of the National Institutes of Health.

### Competing Interest Statement

The authors have declared no competing interest.

